# Age-corrected model for predicting pupil diameter in real-world conditions from melanopic equivalent daylight illuminance

**DOI:** 10.64898/2026.08.05.742771

**Authors:** Manuel Spitschan

**Author notes:** **Corresponding author:** Manuel Spitschan, PhD.

## Abstract

**Purpose:** Pupil diameter in daily life depends on both the light reaching the eye and the observer’s age, but established prediction formulas require laboratory quantities that are rarely measured in natural environments. We developed a compact age-corrected model that predicts pupil diameter from melanopic equivalent daylight illuminance (mEDI).

**Methods:** We used an existing field dataset in which binocular pupil diameter and near-corneal spectral irradiance were recorded while 83 adults aged 18–87 years moved through indoor and outdoor environments. The analysis included 10,082 valid paired observations. We fitted a bounded sigmoid relating pupil diameter to mEDI and age, with each participant given equal influence, and assessed prediction in participants excluded from model fitting. Performance was compared with simpler models, a flexible generalised additive model (GAM), and Watson–Yellott predictions based on assumed field geometry.

**Results:** Pupil diameter decreased smoothly as mEDI increased. Age primarily reduced the difference between pupils in dim and bright conditions, by 0.768 mm per decade, while the predicted bright-light diameter changed little with age. In held-out participants, the bounded model had a participant-balanced root mean squared error (RMSE) of 0.630 mm and mean absolute error of 0.537 mm. The GAM had a slightly lower point-estimate RMSE of 0.610 mm, but the difference was small and uncertain. The bounded model outperformed the tested log-linear, reduced, age-only, and Watson–Yellott alternatives.

**Conclusion:** Age and mEDI are sufficient to provide useful population-average pupil predictions across the observed adult age and real-world light range. The model is transparent, physiologically bounded, and nearly as accurate as a flexible GAM, but predictions approaching darkness remain uncertain because valid mEDI measurements were not available in that range.

**Key points:**

- A compact equation predicts population-average pupil diameter from age and mEDI alone.
- Age mainly compresses the pupil’s response range by reducing pupil diameter under dimmer conditions.
- Prediction error in unseen participants was close to that of a flexible GAM, without requiring a fitted smooth object.
- The model is intended for the observed adult age and field-light range, not for extrapolation into darkness.

## Introduction

Pupil diameter determines retinal illuminance and influences optical image quality. It varies strongly with ambient light and also becomes smaller with age. Existing population formulas capture these relationships under controlled conditions, most notably the Watson–Yellott equation, which predicts steady-state pupil size from photopic luminance, adapting-field area, age, and viewing configuration [1]. These inputs are appropriate for defined laboratory fields but are often unavailable in daily life, where the light at the eye changes continuously and the surrounding scene is neither uniform nor fixed.

Natural-light measurements offer a more direct route to prediction. Melanopsin-expressing retinal ganglion cells contribute to the pupil light response, and mEDI provides a standardised measure of melanopic stimulation [2]. In an earlier field study, wearable binocular pupillometry was combined with near-corneal spectroradiometry as adults moved through indoor and outdoor settings. Pupil diameter was strongly related to light exposure and age, with melanopic quantities explaining the observed variation better than photopic quantities [3]. That study established the empirical relationship but did not provide a portable equation for estimating pupil diameter from readily available inputs.

We therefore asked whether the field relationship could be reduced to a practical age-corrected model. The aim was not to identify the retinal mechanism of pupil control, but to predict population-average pupil diameter using only age and mEDI. The model needed to generalise to people not used for fitting, preserve the expected monotonic decrease in pupil size as light increases, and avoid impossible predictions. We compared this constrained equation with both simpler alternatives and a flexible statistical benchmark to determine how much accuracy was gained or lost by choosing portability.

## Methods

### Data source and analysis sample

We analysed the public dataset from the natural-vision study by Lazar and colleagues [3]. In that study, adults wore a binocular video-based pupillometer and a spectroradiometer that measured spectral irradiance near the corneal plane while they moved through everyday indoor and outdoor environments. Applying the source study’s documented quality and participant-retention criteria produced 10,082 valid pupil-light pairs from 83 adults aged 18–87 years, spanning mEDI values from 1.02 to 58,559 lx. The reconstructed dataset reproduced the source participant counts, key variables, cohorts, and published age regressions in all 19 prespecified checks.

Sampling density differed between participants because their light exposure changed naturally over time. To prevent people with more observations from dominating the model, we calculated each participant’s median pupil diameter within five decade-wide mEDI ranges and gave every participant equal total weight. Measurements from the labelled dark phase were not treated as zero mEDI because they fell below the valid range of the light instrument.

### Prediction model

The model describes a smooth transition between a larger pupil at low mEDI and a smaller pupil at high mEDI:

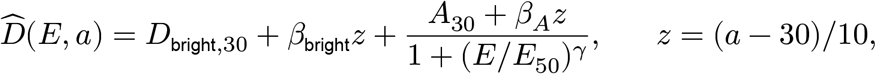

where 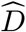 is pupil diameter in millimetres, *E* is mEDI in lux, and *a* is age in years. *D*_bright,30_ is the bright-light pupil diameter at age 30, and *A*_30_ is the difference between low-light and bright-light pupil diameter at that age. *β*_bright_ and *β*_*A*_ describe their change per decade, while *E*_50_ and *γ* determine the position and shape of the light response. The fit was constrained so that pupil diameter remained between 1 and 9 mm across the observed ages and could not increase as mEDI increased.

### Model evaluation

Prediction was assessed with ten repeats of participant-grouped 10-fold cross-validation. Each participant appeared in either the training or test data within a fold, never both. Error estimates were balanced across participants and summarised using RMSE and mean absolute error (MAE). We compared the full model with a reduced bounded equation, a log-linear age-by-light model, age-only and intercept-only baselines, and a GAM that served as a flexible benchmark. We also calculated Watson–Yellott predictions using incident photopic illuminance and an assumed uniform 60-degree adapting field. This comparison was necessarily scenario-based because adapting luminance and field size were not measured in the source study.

The final coefficients were estimated from all participant-by-light-range summaries, with uncertainty obtained from 1,000 participant bootstrap samples. Sensitivity analyses varied data-retention thresholds, light metrics, fitting resolution, treatment of below-detection measurements, and the assumed field geometry for Watson– Yellott predictions. These analyses were used to distinguish stable predictive findings from parameters that depended on how the low-light data were represented.

## Results

### Light and age shape different parts of the pupil response

Across the observed field range, pupil diameter decreased continuously as mEDI increased (Figure 1). The effect of age was most evident at lower light levels. At age 30, the model estimated a bright-light diameter of 1.621 mm and a further low-light expansion of 6.488 mm. This expansion decreased by 0.768 mm per decade, whereas the age-related change in the bright-light asymptote was small and compatible with no change. The model therefore describes ageing primarily as a compression of the pupil’s operating range rather than a uniform downward shift at every light level.

**Figure 1.**
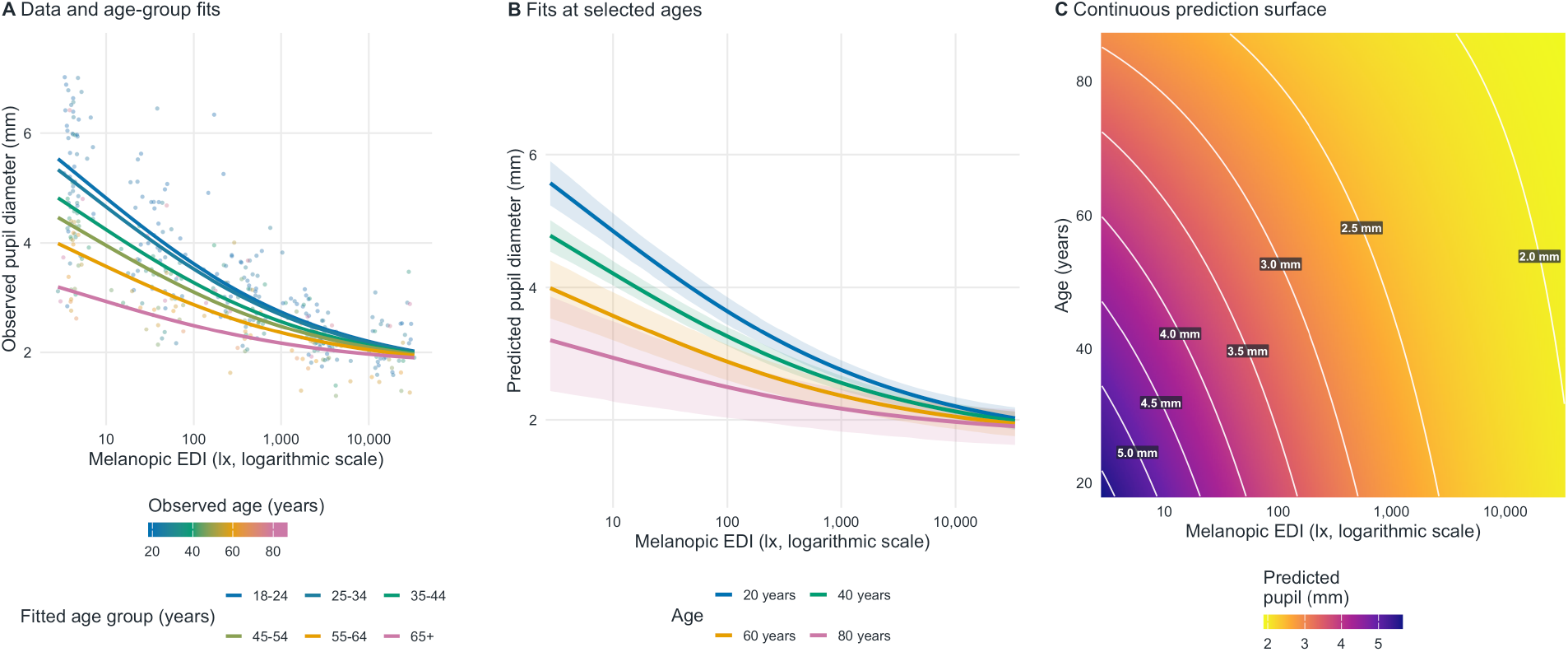
Observed and predicted relationships between pupil diameter, mEDI, and age. Panel A shows participant-specific median pupil diameters and fitted curves for the source study’s age groups. Panel B shows predicted curves at ages 20, 40, 60, and 80 years with pointwise 95% participant-bootstrap intervals. Panel C shows the continuous prediction surface across the observed age and mEDI range.

The estimated response midpoint was 5.19 lx. Its uncertainty was substantially greater than that of the age effect because the source study did not contain valid continuous mEDI measurements in darkness. The data clearly supported the overall decline in pupil diameter across the measured range, but provided less information about the exact position of the low-light asymptote.

**Table 1:** Model coefficients and prediction performance. Coefficient intervals are participant-bootstrap intervals; performance was measured in held-out participants and balanced across participants.

| Section | Parameter or model | Estimate or performance |
| --- | --- | --- |
| Coefficient | Bright-light diameter at age 30 | 1.621 (1.172 to 1.922) |
| Coefficient | Additional low-light diameter at age 30 | 6.488 (5.341 to 6.921) |
| Coefficient | Bright-light change per decade | 0.025 (-0.039 to 0.087) |
| Coefficient | Low-to-bright response-range change per decade | -0.768 (-0.972 to -0.517) |
| Coefficient | Response midpoint, mEDI | 5.185 (3.078 to 18.241) |
| Coefficient | Response-shape exponent | 0.316 (0.261 to 0.389) |
| Held-out prediction | Age-corrected bounded model | RMSE 0.630 mm; MAE 0.537 mm |
| Held-out prediction | GAM | RMSE 0.610 mm; MAE 0.515 mm |
| Held-out prediction | Log-linear model | RMSE 0.653 mm; MAE 0.553 mm |

| Section | Parameter or model | Estimate or performance |
| --- | --- | --- |
| Held-out prediction | Reduced bounded model | RMSE 0.663 mm; MAE 0.559 mm |
| Held-out prediction | Watson–Yellott, assumed 60-degree field | RMSE 0.836 mm; MAE 0.689 mm |
| Held-out prediction | Age only | RMSE 1.148 mm; MAE 0.971 mm |
| Held-out prediction | Intercept only | RMSE 1.185 mm; MAE 0.994 mm |

### A portable model retained nearly all predictive accuracy

For participants excluded from fitting, the bounded model achieved a participant-balanced RMSE of 0.630 mm and an MAE of 0.537 mm. The GAM had the lowest point-estimate RMSE at 0.610 mm, but its advantage over the bounded model was only 0.020 mm and the 95% interval included no difference (−0.052 to 0.012 mm). The added flexibility of the GAM therefore produced little demonstrable improvement in prediction.

The bounded model performed better than the log-linear model by 0.022 mm and better than the reduced bounded model by 0.033 mm. Models based on age alone or a single population mean performed much worse, confirming that current light exposure carried essential predictive information. The assumed 60-degree Watson–Yellott scenario had an RMSE of 0.836 mm, compared with 0.630 mm for the new model. None of the held-out predictions from the bounded model fell outside the prespecified physiological range of 1–9 mm.

### The age effect was stable, but darkness remained weakly constrained

The reduction in response range with age was consistent across the sensitivity analyses. By contrast, the estimated response midpoint changed when the temporal or mEDI-bin resolution was altered. Watson–Yellott performance also depended on the assumed adapting-field geometry, with RMSE ranging from 0.819 to 0.889 mm across tested field sizes. These findings point to the same boundary: the data support prediction across observed real-world light levels more strongly than they support inference about an unmeasured dark asymptote.

## Discussion

This study converts a rich field dataset into a practical equation for predicting pupil diameter from two inputs that can be measured or specified directly: mEDI and age. The central pattern is straightforward. Pupils became smaller as melanopic light exposure increased, and age reduced the size of this light-dependent change. Older and younger adults differed most under dimmer conditions, while their predicted pupil diameters converged under bright conditions. This representation is more informative than treating age as a constant correction applied equally at all light levels.

The model’s main contribution is portability rather than maximal statistical flexibility. A GAM achieved a marginally lower point-estimate error, but the difference was small and uncertain. The bounded equation retained nearly all of that predictive performance while providing an explicit six-parameter formula, a monotonic light response, and predictions constrained to a plausible pupil range. It can therefore be implemented in software, optical calculations, study planning, or light-exposure analyses without distributing a fitted statistical object. Its improvement over age-only models also shows why a population-average pupil value is inadequate when light exposure varies substantially.

Comparison with Watson–Yellott should be interpreted narrowly. The classical equation predicts pupil diameter from adapting luminance and field geometry, whereas the field dataset measured incident illuminance at the eye and did not characterise a uniform adapting field. The scenario analysis therefore does not show that the new model supersedes Watson–Yellott under its intended laboratory conditions. It shows that an mEDI-based model is more useful when the available measurements come from wearable light sensors in uncontrolled environments and the classical geometric inputs are unknown.

The most important limitation concerns very low light. Measurements labelled as dark were below the spectroradiometer’s valid range, so the model was not anchored by measured mEDI values approaching zero. Predictions across the observed field range were stable, but the exact low-light asymptote and response midpoint depended more on modelling choices. The model should consequently be used for interpolation within the observed range of adults aged 18–87 years and mEDI values of approximately 1.02 to 58,559 lx. It requires independent validation before application to children, clinical populations, different measurement systems, or controlled conditions outside that range. A measured pupil diameter remains preferable whenever an individual measurement is available.

For practical use, the equation requires no participant-specific calibration. The [Melanopic Pupil](https://tscnlab.shinyapps.io/MelanopicPupilPredictor/) Predictor provides an interactive implementation, and the coefficients can be incorporated directly into other tools. Future field studies that include calibrated measurements in darkness and direct estimates of adapting-field geometry would strengthen low-light predictions and allow a more direct comparison with luminance-based formulas.

## Appendix Expanded methodological protocol and supporting analyses

### A1. Source-study participants, eligibility, and recording protocol

The source was our openly released, pseudonymised dataset from the registered-report field study of Lazar and colleagues [3]. Recruitment was organised across six intended age groups: 18–24, 25–34, 35–44, 45–54, 55–65, and older than 65 years. Healthy community volunteers were eligible if they did not require refractive correction and did not wear contact lenses during measurement, had visual acuity of at least 20/40 on a Snellen chart, had normal colour vision on the Hardy–Rand–Rittler test, and passed screening for relevant ocular, neurological, metabolic, medication, mental-health, drug, and alcohol factors. The source report records 113 people invited to screening, 106 assigned to the protocol, 87 completing technically usable trials, and 83 remaining after the adjusted participant-level data-loss criterion used for the primary reconstruction. Ethical approval for the source study was granted by the Ethikkommission Nordwest- und Zentralschweiz (project 2019-01832) [3].

Each participant completed an approximately one-hour ambulatory protocol containing natural indoor and outdoor tasks, a 10 min dark-adaptation period, and a 14 min controlled laboratory-light period. Pupil images and spectral irradiance were acquired synchronously at intervals of 10 s, producing approximately 300– 360 scheduled samples per participant. Natural conditions included fluorescent and LED room lighting, computer-screen light, window-transmitted daylight, and outdoor daylight. The present primary model used only observations adjudicated in the released data as field measurements. Dark and laboratory observations were retained for reconstruction checks and explicitly labelled sensitivity analyses, but were not silently combined with the primary field observations.

Pupil images were acquired with an infrared Pupil Labs video eye tracker. Before ambulatory recording, a 3 min calibration was used to fit and visually verify the manufacturer’s three-dimensional eye model across head and eye positions. Absolute pupil diameter in millimetres was subsequently estimated with Pupil Labs software version 1.15.71. The released outcome used here was the source variable diameter_3d, which is based on the segmented elliptical pupil contour and the fitted rotating-eye model.

Spectral irradiance was recorded with an STS-VIS spectroradiometer (Ocean Insight, Oxford, UK) fitted with a CC-3-DA cosine corrector. The source instrument covered 350–800 nm, with reported wavelength accuracy of +/-0.13 nm, 6 nm optical resolution, a signal-to-noise ratio greater than 1500:1, and a 4600:1 dynamic range. It was mounted centrally on the forehead, approximately coplanar with the cornea, directed 15 degrees below horizontal, and sampled a 180-degree field. The measurement is therefore a near-corneal-plane proxy for incident ocular light, not a direct measurement of retinal irradiance or the luminance distribution of the viewed scene.

### A2. Data reconstruction, spectral quantities, exclusions, and analysis unit

The reconstruction began with all 113 participant directories and 32,810 scheduled sampling rows in the source repository. Participant files, the cleaned survey object, manually adjudicated phase labels, and corrected spectral calculations were treated as separate source components. Participant and within-participant sample identifiers were used for all joins; row order was never used as an implicit key. Processed source objects were used only for information that could not be recovered from participant files, particularly the final field labels and the mapping between pupil samples and corrected spectral records.

The source spectra had been resampled by energy-preserving interpolation to 1 nm spacing from 380 to 780 nm. Melanopic irradiance and melanopic equivalent daylight illuminance (mEDI) were calculated using the CIE S 026 melanopsin spectral sensitivity, and photopic illuminance was calculated from the CIE photopic luminous-efficiency function [2]. The present analysis used the corrected mEDI supplied by the source spectral-processing workflow.

We reapplied the source quality rules to the participant files. Pupil estimates with Pupil Labs confidence below 0.60 were set to missing, as were diameters outside the prespecified physiological interval of 1–9 mm. Recorded light values equal to zero were treated as invalid. Participant data loss was calculated as the proportion of scheduled samples lacking either a valid pupil estimate or a valid melanopic light measurement.

Participants with more than 75% data loss were excluded from the primary cohort, reproducing the source study’s adjusted full-cohort rule. The original 50% threshold was retained as a sensitivity analysis. No missing pupil, age, or light value was imputed.

Primary analysis rows additionally required inclusion in the source cohort, a manually adjudicated Field label, pupil confidence of at least 0.60, and finite positive mEDI, age, and pupil diameter. These rules yielded 10,082 complete pupil-light observations from 83 adults aged 18–87 years, with corrected field mEDI from 1.024 to 58,559.35 lx. Dark-phase measurements were not assigned an mEDI of zero because both melanopic and photopic measurements below 1 lx lay outside the valid range of the source workflow. Two sensitivity analyses instead assigned explicit boundary values of 0.1 or 0.5 lx to test the dependence of the low-light parameters on this assumption.

The approximately 10 s field series contained unequal numbers of observations per participant and substantial short-lag serial dependence. The primary fitting unit was therefore a participant-by-light-bin median rather than an individual time sample. For each participant, median mEDI, photopic illuminance, and pupil diameter were calculated in five mEDI intervals: [1,10), [10,100), [100,1000), [1000,10,000), and at least 10,000 lx. This produced 376 participant-by-bin summaries. If participant *p* contributed *n*_*p*_ populated bins, each summary received weight *w*_*pi*_ = 1/*n*_*p*_; consequently every participant contributed total weight one, irrespective of recording duration or number of populated light bins. Medians reduced the effect of brief artefacts while retaining the order-of-magnitude exposure structure that motivated the logarithmic light analyses.

Before model fitting, the reconstructed data were checked against the published cohort flow, condition counts, demographics, pupil summaries, corrected-light summaries, and the age regressions reported in Figure 8 of the source article. All 19 formal replication checks passed, including matched participant/sample values in the released processed object. These checks establish that the present analysis operated on the intended source records; they do not constitute external validation of the new prediction equation.

### A3. Development and selection of the prediction equation

Equation development began with four deployment requirements. The prediction had to use only age and mEDI; approach interpretable dim- and bright-light limits; decrease monotonically as light increased; and remain within a plausible pupil range. A Hill-type transition was selected as the simplest familiar saturating function that met those requirements:

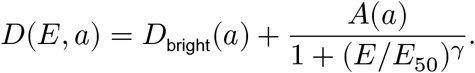

This representation separates the limiting bright-light diameter *D*_bright_, the additional dim-to-bright response amplitude *A*, the half-amplitude exposure *E*_50_, and the transition exponent *γ*. It is preferable for prediction across several orders of magnitude of positive exposure to an unconstrained polynomial or straight line, either of which can diverge outside the centre of the observed range.

The source results and earlier laboratory observations indicate that age differences are greatest in dim light and diminish as illumination increases [3, 4]. A single additive age term cannot represent that pattern. We therefore allowed age to change both the bright-light limit and the response amplitude:

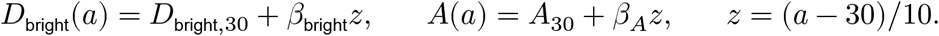

Age was centred at 30 years and expressed in decades so that the two age coefficients have direct units of millimetres per decade and the optimization is numerically well scaled. The midpoint and shape exponent were kept age-invariant. Allowing further age interactions in these weakly constrained parameters would add complexity that was not supported by measurements near darkness. With *A*(*a*) *≥* 0, *E*_50_ > 0, and *γ* > 0, the derivative with respect to mEDI is non-positive, so the fitted pupil cannot dilate as light increases.

The six-parameter equation was the smallest candidate considered that allowed the empirically established age effect to vary with light level while remaining a portable closed form. It was not selected by an unrestricted automated search over functional forms. Because its structure was developed with knowledge of the source pattern, the grouped cross-validation assesses coefficient estimation for this fixed form; it is internal validation and not independent validation of the complete model-development process.

All candidate models used the same participant-bin summaries and weights. The reduced bounded model retained the Hill transition but used an age-invariant amplitude plus one additive age coefficient; it tested whether a light-dependent age effect justified the sixth parameter. The log-linear model,

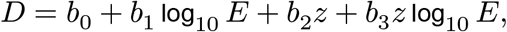

matched the approximately linear log-light analysis in the source paper but did not enforce asymptotes or physiological bounds. Age-only and intercept-only regressions quantified the prediction gained from current light exposure and age. A generalised additive model used penalised smooths of log_10_ *E* and age, with basis dimension 5 for each main effect and a tensor-product interaction with basis dimensions 4 by 4; smoothing was estimated by restricted maximum likelihood. The GAM was retained as a deliberately flexible accuracy benchmark but was not selected as the deployable equation because it requires a stored fitted object, provides less direct parameter interpretation, and does not itself enforce monotonicity or the 1–9 mm range. The Watson–Yellott model was treated as an external scenario formula rather than another fit to the outcome data because its required adapting luminance and field geometry were not measured.

### A4. Estimation, constraints, internal validation, and uncertainty

Coefficients were estimated by minimising participant-weighted mean squared error,

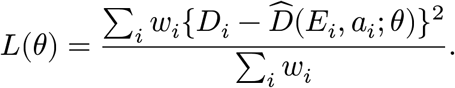

*A*_30_, *E*_50_, and *γ* were optimized on logarithmic scales to enforce positivity. Search bounds were 1–7 mm for *D*_bright,30_, 0.01–8 mm for *A*_30_, −1.5–1.5 mm per decade for each age coefficient, 0.01–10,000,000 lx for *E*_50_, and 0.03–8 for *γ*. Multiple dispersed starting values were fitted with bounded L-BFGS-B optimization: eight starts for the final fit and four starts in each cross-validation training set. The best solution was refined by sequential least-squares quadratic programming using numerical gradients and Jacobians. Explicit inequalities required non-negative response amplitude and both the bright and dim asymptotes to remain within 1–9 mm at the minimum and maximum ages in each training set. Fits were accepted only after optimizer convergence with maximum inequality violation no greater than 10^−7^.

Internal validation used ten repeats of participant-grouped 10-fold cross-validation. Participants were sorted into age-quartile strata and randomly assigned approximately evenly to folds using fixed random-number seeds beginning at 20260804. Within each repetition and fold, every summary from a participant occurred exclusively in the training or test set, and every model component was re-estimated using the training participants before predictions were made for held-out participants. Each participant-bin summary consequently received ten out-of-fold predictions per model. Fold assignments and every held-out prediction were saved.

The primary RMSE was first calculated separately for each participant and then averaged across participants; participant-balanced MAE was calculated analogously. Row-level RMSE and MAE, calibration intercept and slope from regressing observed on predicted diameter, and the fraction of predictions outside 1–9 mm were retained as secondary summaries. Pairwise model comparisons used the within-participant difference in RMSE. Uncertainty in each mean paired difference was estimated using 2,000 nonparametric bootstrap resamples of participants.

After internal validation, the reported coefficients were refitted to all 376 participant-bin summaries. Parameter uncertainty used 1,000 nonparametric participant bootstrap samples. All summaries for a sampled participant were retained together; when a participant was selected more than once, duplicate selections were assigned distinct bootstrap identifiers before refitting. Reported 95% intervals are the 2.5th and 97.5th percentiles of successful fits. Sensitivity analyses repeated the fit with the 50% participant-loss cohort, a pupil-confidence threshold of 0.70, participant-balanced row-level data, uncorrected mEDI, melanopic irradiance, photopic illuminance, half-decade light bins, assumed dark boundaries of 0.1 and 0.5 lx, removal of the highest-error participant, and alternative age-reference parameterisation.

### A5. Watson–Yellott scenario comparison

The Watson–Yellott model predicts pupil diameter from adapting luminance, angular field area, age, and monocular or binocular viewing [1]. It is a luminance formula, not an irradiance or illuminance formula. Because the source field study measured incident photopic illuminance at the forehead and did not recover the spatial luminance distribution of the viewed scene, an exact Watson–Yellott prediction cannot be calculated from these records. The comparison therefore imposed a reproducible geometric scenario. For a centred uniform circular field of diameter *d*, incident photopic illuminance *E*_*V*_ was converted to field luminance as

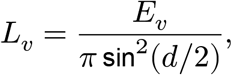

where *d*/2 is expressed in radians. Watson–Yellott then used angular area *π*(*d*/2)^2^ in square degrees, continuous age, and binocular viewing. Stanley and Davies measured diameters from 0.4 to 25.4 degrees and cautioned against extrapolation beyond that range. Watson and Yellott nevertheless state that they did not restrict field diameter because Bouma’s results suggested integration over much larger fields, and their article illustrates the unified formula at 60 degrees [1]. Figure S1 therefore uses *d* = 120 degrees as a broad full-field scenario and the largest diameter in the systematic sensitivity analysis. This remains an extrapolation based on an idealised centred uniform field, not measured scene geometry. The Watson implementation was also checked numerically against both the source-transcribed MATLAB implementation and the published unified form over a grid of luminances, ages, field sizes, and viewing modes.

**Figure S1.**
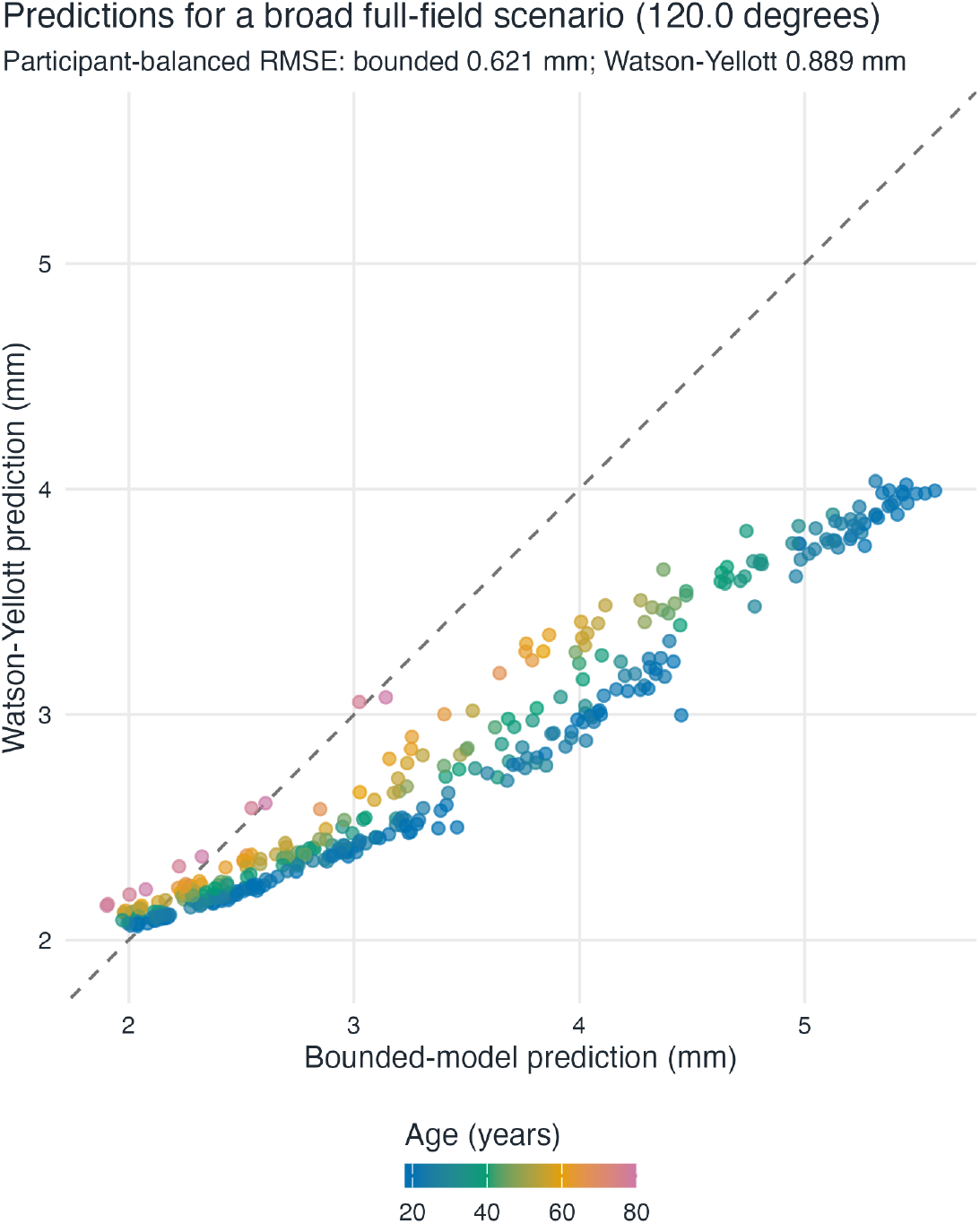
Direct comparison of participant-held-out bounded-model and Watson–Yellott predictions for an idealised 120-degree centred uniform field, used as a broad full-field scenario. This plot was Figure 6 in the supplementary-analysis report and is relabelled S1 here. The dashed line is identity and points are coloured continuously by age. The comparison includes 366 participant-bin summaries from 81 participants with valid photopic illuminance and age within the verified Watson–Yellott range.

At 120 degrees, participant-balanced RMSE was 0.621 mm for the bounded model and 0.889 mm for Watson–Yellott. The mean within-participant RMSE disadvantage for Watson–Yellott was 0.268 mm, with a participant-bootstrap 95% interval of 0.165 to 0.363 mm. The two predictions remained strongly associated, but Watson–Yellott tended to give smaller values where the bounded model predicted larger pupils. This result shows the consequence of applying a fully specified broad-field assumption to field-sensor data. It does not demonstrate that the new equation supersedes Watson–Yellott under controlled conditions where adapting luminance and field geometry are actually measured.

### A6. Relationship between photopic illuminance and mEDI

**Figure S2.**
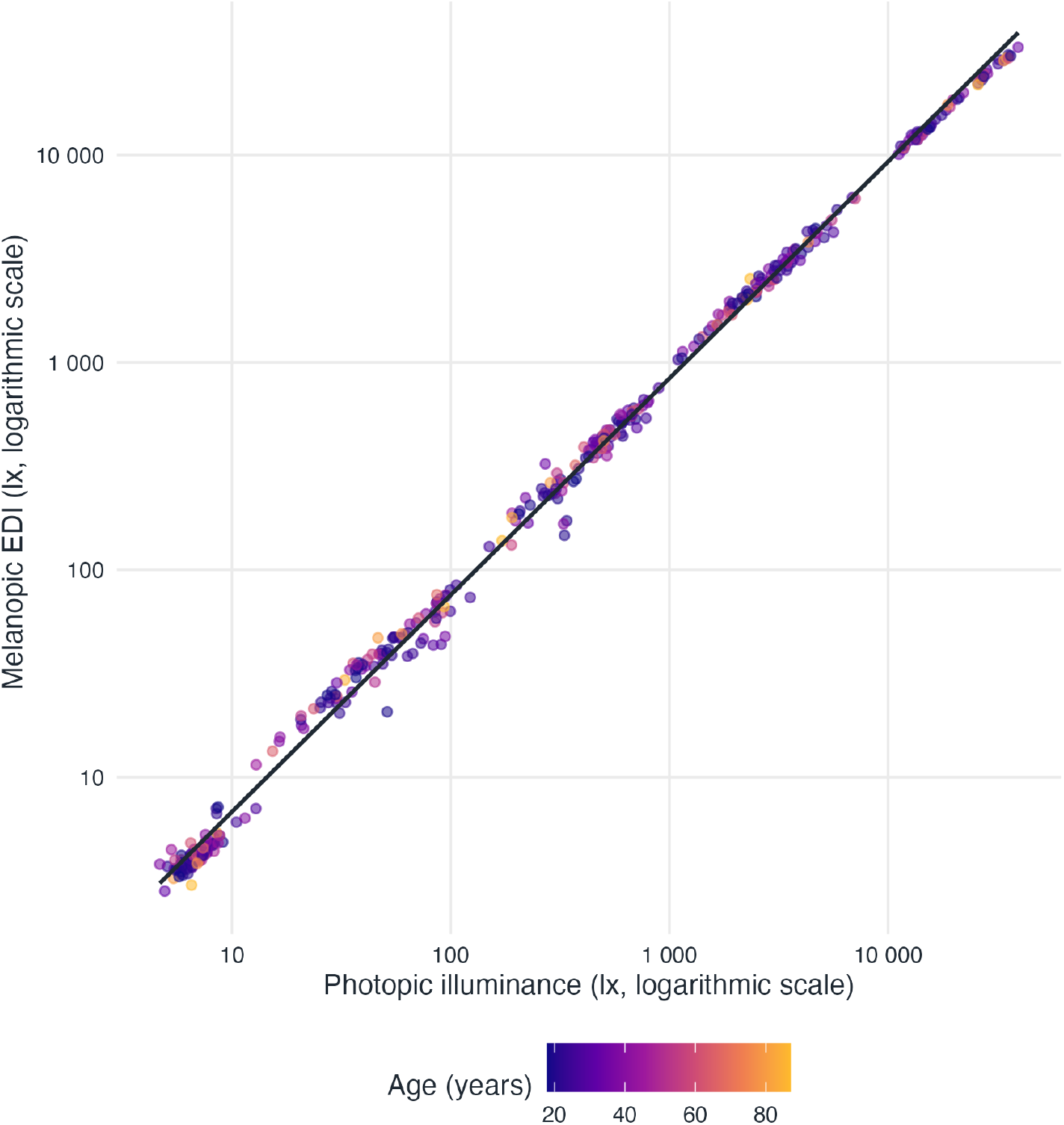
Relationship between photopic illuminance and melanopic EDI in the 376 participant-by-light-bin summaries. This plot was Figure 8 in the supplementary-analysis report and is relabelled S2 here. Both axes are logarithmic, points are coloured continuously by age, and the solid line is a descriptive log-log linear fit.

Photopic illuminance and mEDI were necessarily related because both quantities were calculated from the same recorded spectra and both increase with overall light level. Across the participant-bin summaries, their log-scale correlation was 0.9988 and the descriptive log-log regression had *R*^2^ = 0.9975. This near-linear relationship explains why changing the predictor from mEDI to photopic illuminance did not remove the broad light-response pattern. It does not establish physiological interchangeability: the ratio between photopic and melanopic quantities depends on spectral composition, and the present natural-light sample may contain less independent spectral variation than other lighting environments. The headline equation therefore uses mEDI because the source study found melanopsin-weighted quantities to provide the stronger single-predictor description and because mEDI is defined by a standardised spectral weighting [2, 3].

### A7. Calibration of participant-held-out predictions

**Figure S3.**
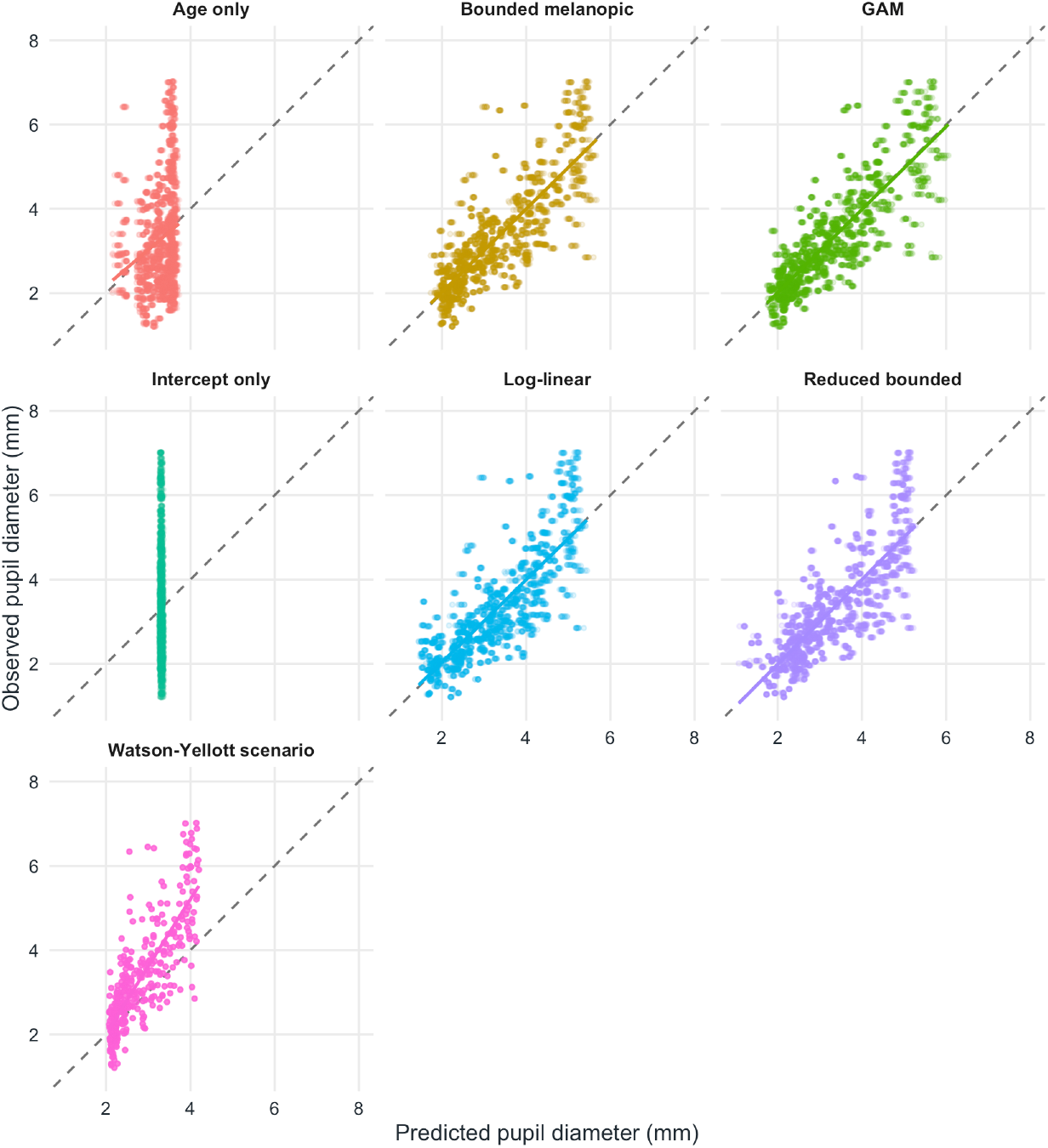
Calibration of participant-held-out predictions across ten repeats of grouped 10-fold crossvalidation. This plot was Figure 9 in the supplementary-analysis report and is relabelled S3 here. Each point represents an out-of-fold prediction for a participant-bin summary; solid lines are model-specific linear calibration fits and dashed lines indicate identity.

The bounded melanopic model lay close to identity, with calibration intercept 0.042 mm and slope 0.988. The GAM was similarly calibrated, with intercept 0.042 mm and slope 0.985. The scatter around both fitted lines shows that these are population-average rather than participant-specific predictions. Age-only and intercept-only models lacked the exposure-dependent range visible in the observations, while the Watson– Yellott scenario showed stronger calibration deviation under its assumed 60-degree field. Together with the participant-balanced error measures, this figure shows that the main conclusion is not based only on in-sample fit: the bounded equation preserved the observed scale and light-dependent range in participants excluded from coefficient estimation.

### A8. Which Watson–Yellott field size best matches the new predictions?

We additionally asked whether one assumed Watson–Yellott field diameter could make its predictions closely correspond to the bounded model’s predictions for the real-world field data. For every diameter in the systematic 0.4–120-degree grid, we calculated the root mean squared difference between Watson–Yellott and bounded-model predictions separately within each participant and then averaged those participant-specific values. This participant-balanced prediction RMSD compares the two prediction systems directly; it is distinct from RMSE against observed pupil diameter.

**Figure S4.**
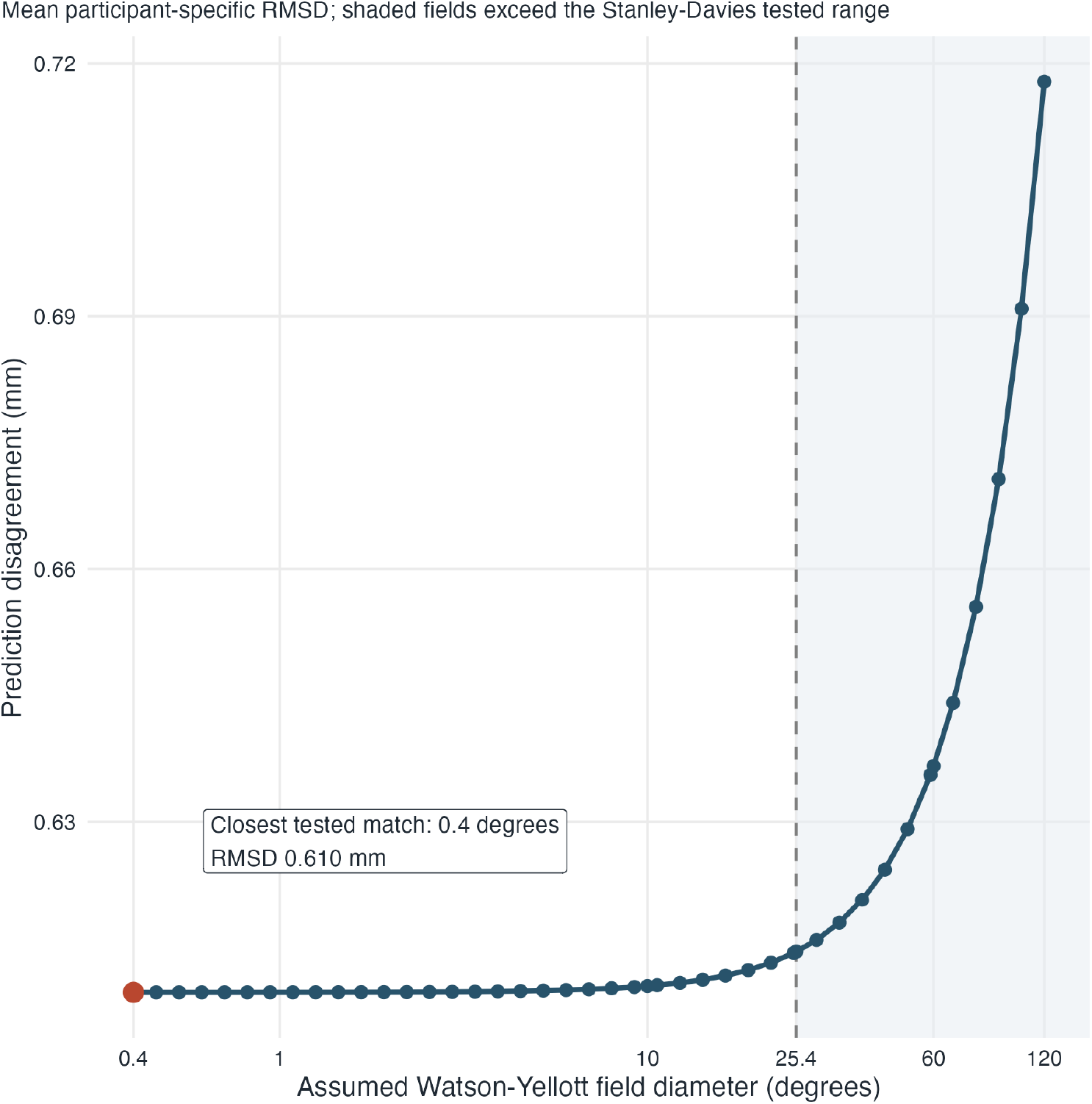
Disagreement between participant-held-out bounded-model and Watson–Yellott predictions across assumed circular field diameters. Lower values indicate closer correspondence. The highlighted point is the closest match on the tested grid. The shaded region exceeds the 0.4–25.4-degree field-size range measured by Stanley and Davies, although Watson and Yellott did not impose that range as a formula limit.

The closest tested match occurred at the lower grid boundary of 0.4 degrees, with participant-balanced prediction RMSD 0.610 mm. Disagreement was 0.615 mm at 25.4 degrees and increased to 0.718 mm in the 120-degree broad full-field scenario. Because the curve is nearly flat across the smallest diameters and its minimum occurs at the search boundary, the data do not identify a unique interior “equivalent field”. Most importantly, the 0.4-degree result must not be interpreted as the physical field of the real-world observations. It is only the effective Watson–Yellott setting that minimises prediction disagreement under the imposed uniform-field conversion. Field size alone cannot reconcile the remaining difference between a luminance-and-geometry formula and a melanopic field model.

### A9. Computational reproducibility

Analyses were run in R 4.5.0 and the manuscript was rendered with Quarto 1.6.43. Package versions are recorded in renv.lock; fixed random-number seeds, fold assignments, bootstrap estimates, derived tables, and fitted objects are retained with the code. From the repository root, source(“scripts/run_all.R”) reconstructs the data, repeats all model fits and validation analyses, regenerates figures and tables, runs the automated tests, and renders the HTML and PDF reports. The test suite checks reconstruction counts, source-result replication, model monotonicity and physiological bounds, participant-grouped validation, invariance to age-reference reparameterisation, calculator consistency, and the Watson–Yellott translation.

The analysis repository is https://github.com/tscnlab/Spitschan_OphthalmicPhysiolOpt_2026. The source-study code and data are available at https://github.com/tscnlab/LazarEtAl_RSocOpenSci_2024. The source study also deposited processed data at https://doi.org/10.6084/m9.figshare.24230848.v3 and supporting material at https://doi.org/10.6084/m9.figshare.24230890.v1.

## Declarations

### Ethics approval and consent to participate

No new data were collected for this analysis. The original study was approved by the cantonal ethics commission, Ethikkommission Nordwest- und Zentralschweiz (project 2019-01832), as reported in the source publication [3].

### Consent for publication

Not applicable.

## Author contributions

M.S.: Conceptualization, Data curation, Funding acquisition, Methodology, Project administration, Resources, Validation, Visualization, Writing - Original draft, and Writing - Review and editing.

## Data and code availability

All data and code needed to reproduce the analysis and manuscript are available at https://github.com/tscnlab/Spitschan_OphthalmicPhysiolOpt_2026. The underlying field dataset was originally reported by Lazar and colleagues [3]. The interactive Shiny calculator is available as the [Melanopic Pupil Predictor](https://tscnlab.shinyapps.io/MelanopicPupilPredictor/).

